# Microbes as manipulators of developmental life-history

**DOI:** 10.1101/2024.02.02.578589

**Authors:** Matthew C. Kustra, Tyler J. Carrier

## Abstract

Marine invertebrates mainly reproduce by energy-poor eggs that develop into feeding larvae or energy-rich eggs that develop into non-feeding larvae^1–4^. Transitions between these reproductive strategies have been studied in detail^5–7^, yet the evolutionary factor(s) responsible for these switches remains elusive. Here, we use theoretical models to show that microbes with the capacity to manipulate host reproduction are one possible factor. We report that microbial manipulators create a sperm-limited environment that selects for larger eggs by shifting the host’s sex ratio towards female dominance and, as a result, serve as the evolutionary driver of transitions in the developmental life-history for marine invertebrates. Loss of a microbial manipulator can then recover the ancestral developmental life-history. We also document more than a dozen genera of marine invertebrates from throughout the world’s oceans that fit the framework of a microbe-induced switch between these predominate reproductive strategies. We anticipate that microbial manipulators have a yet-to-be appreciated influence on the life-history strategies of marine invertebrates. We find it paramount to understand if transitions in developmental life-history also occur without microbial manipulators as well as if the underlying mechanisms of these manipulations are convergent with terrestrial systems.

## MAIN

The most common reproductive strategy amongst marine invertebrates involves producing a high number of small, energy-poor eggs that develop into feeding larvae (*i.e.*, planktotrophy)^2,8–10^. This stability has occasionally been disrupted over the last several hundred million years by a rapid switch in developmental life-history^2,8–10^. Species with the derived reproductive strategy produce fewer large, energy-rich eggs and larvae that undergo metamorphosis without feeding (*i.e.*, lecithotrophy)^2,8–10^. Transitions in developmental and reproductive life-history have occurred in most major animal phyla, are primarily unidirectional, and intermediates between planktotrophy and lecithotrophy are rare^8–11^. Despite the patterns of transitions in developmental life-history having been studied in detail^4–7,12–15^, the factor(s) responsible for driving switches in reproductive strategy remain elusive.

Microbes may be one such factor, as they are functionally beneficial to animal reproduction^16^ and have evolved mechanisms to override components of this host program^17,18^. Microbial manipulators are often inherited through the cytoplasm of the egg and manipulate the host to favor their transmission^17,18^. Hosts affected by reproductive manipulators (*e.g.*, *Wolbachia*) are diverse and are particularly common in terrestrial arthropods and nematodes, where >60% of species are thought to be infected^17–19^. Microbes that are phylogenetically related to terrestrial reproductive manipulators are increasingly found in the ocean and, specifically, in association with major marine invertebrate phyla (*e.g.*, the sea urchin *Heliocidaris erythrogramma*)^20–23^. There is considerable overlap in the types of reproductive manipulation and the phylogenetic breadth of potential associations between terrestrial and marine hosts despite their vastly different life-history tendencies^18,22,24^.

The primary difference between reproductive manipulators in these two environments is that there is an apparent unbalanced influence on the two predominant developmental life-histories found in the ocean^18,22^. Microbes that manipulate reproduction are thought to have a parasitic effect on the ancestral planktotroph and a mutualistic effect on the derived lecithotroph^22^. This would create an evolutionary gradient where reproductive manipulators could serve as a selective agent to induce transitions between these major reproductive strategies. We tested this by simulating the eco-evolutionary dynamics of egg size (diameter, μm) for free-spawning marine invertebrates following a novel association with a microbial manipulator. We report that microbial manipulators create a sperm-limited environment that selects for larger eggs by shifting the host’s sex ratio towards female dominance and, as a result, serving as the evolutionary driver of transitions in the developmental life-history for marine invertebrates. We also find more than a dozen genera of marine invertebrates that fit the framework of a microbe-induced switch in developmental life-history.

## RESULTS

### Microbial manipulators alter egg size

We used a grid optimum model to test whether egg size evolves following a novel association with a microbe that manipulates host reproduction by feminization (*i.e.*, the conversion of genetic males to females) or male killing (*i.e.*, microbe-induced male mortality) and can compensate host fitness by supplementing host nutrition (*i.e.*, enhanced growth)^18,25^. We simulated the population dynamics of free-spawning marine invertebrates across a range of potential egg size-offspring survival relationships, with the shape of this relationship determined by the *B* parameter (Fig. S1)^3,26^. As such, larger *B* values indicate ecological scenarios that increase the importance of larger egg size for offspring survival (Fig. S1). We assume that all females produce eggs of the same size, that fecundity is inversely proportional to egg size (*i.e.*, maternal investment), and that individuals with larger eggs have an increased probability of fertilization and survival^2,3,8^. Once the populations stabilized, we calculated the fitness of resident (a) and mutant (a ± 10 μm) egg sizes and then record the direction of evolution (Fig. 1A).

**Fig. 1:**
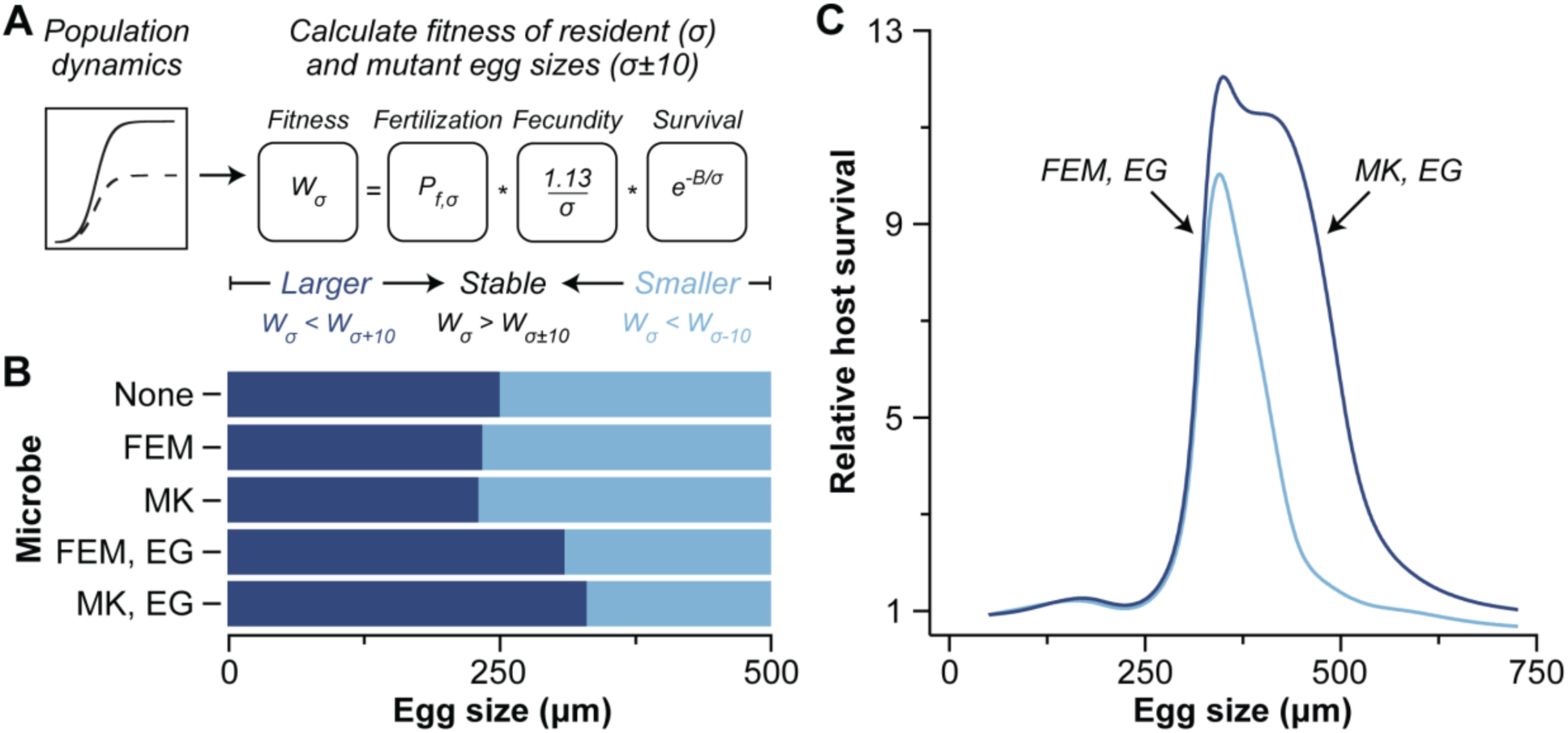
Microbial manipulators alter egg size. (A) An invasion grid approach was used to determine whether egg size evolves following a novel association with a microbe that manipulates host reproduction by feminization (FEM) or male killing (MK) and can compensate host fitness by supplementing host nutrition (EG). The fitness of an individual producing the resident egg size (a) was calculated and compared to rare mutants that produced either a slightly smaller or larger egg sizes (a ± 10 μm). Stable points were identified by the resident egg size having greater fitness than either mutant. (B) Evolutionary direction of egg size with no microbe, FEM (*r =* 0.69), MK (m_k_ =0.69), and FEM or MK plus EG (*g* = 1.2) at a *B* parameter value of 800. (C) Relative host survival following a novel association with a microbe that manipulates by FEM or MK plus EG was determined using the stable egg size across *B* values and back-calculating for survival (*i.e.*, fitness) relative to a population that did not associate with a microbial manipulator.

We observed that the presence of a microbial manipulator altered the egg size of free-spawning marine invertebrates (Fig. 1B). The magnitude and direction of egg size evolution depended on the interaction between manipulation, enhanced growth, and the underlying ecology (*i.e.*, the *B* parameter; Fig. S1-S3). Male killing and feminization alone resulted in a slight decrease in egg size across most ecological scenarios, while the combination of manipulation and enhanced growth led to the evolution of larger eggs (Fig. 1B, S1-S3). The latter, however, was dependent on the underlying ecology. Stable egg size was larger for free-spawning marine invertebrates that associated with a microbe that manipulates by male killing and enhanced growth compared to no manipulation across all *B* values. A similar pattern was observed for a microbe that manipulates by feminization and enhanced growth until a *B* value of >2,600, after which egg size was smaller with a microbial manipulator than without one (Fig. S1-S3).

Evolutionary shifts in stable egg sizes due to the microbial manipulator also led to notable differences in the fitness of populations of free-spawning marine invertebrates (Fig. 1C). Specifically, fitness with and without a microbial manipulator was similar for populations that had egg sizes between 50 and ∼250 μm. Relative fitness then increased to a local maximum of ∼10.0x and ∼12.0x at an egg size of ∼345 μm and ∼350 μm for populations with a manipulator that induced feminization or male killing and enhanced growth, respectively (Fig. 1C). Relative fitness was then similar to baseline at an egg size of ∼520 μm and∼ 665 μm for populations with a manipulator that induced feminization or male killing and enhanced growth, respectively (Fig. 1C). As such, the male killing manipulator had a higher peak (∼1.2x) and broader distribution (∼1.7x) of relative fitness compared to a feminization manipulator. Interestingly, this local maximum in relative fitness coincided with the egg sizes of marine invertebrates that exhibit the derived developmental mode (*i.e.*, lecithotrophy) and, thus, implying that microbes could be the inductive factor along this fitness gradient and of transitions in developmental life-history^1,4,27^.

### Microbial manipulators induce transitions in developmental life-history

The grid optimum model implies that the increase in egg size due to a novel association with a microbial manipulator could induce a transition from the ancestral planktotroph to the derived lecithotroph (Fig. 1). This approach has limitations because it assumes that egg size does not vary within a population and that the underlying ecological and evolutionary processes are separate. This model is also unable to assess the relative speed of these transitions and the sequence of events leading to a switch in developmental life-history. Therefore, we established a quantitative genetic population model that relaxes these assumptions, allowing us to directly test whether a novel association with a microbial manipulator could serve as the selective pressure of egg size evolution that, in turn, induces a transition in developmental life-history.

This quantitative genetic population model assumes that egg size is a normally distributed trait in a population [x-± 10 standard deviation (s.d.) of 9 μm] and calculates the egg size-based fitness of females using the same equations as the invasion grid approach (Fig. 1A). Mean egg size of each generation was calculated using the quantitative genetic breeder’s equation for the surviving adults from the previous generation and their surviving offspring. Constant variation was assumed for each generation, and iterations were performed until the mean egg size reached a stable point. Bacterial abundance (*i.e.*, titer) also scales allometrically with offspring size^28–30^ and, therefore, we established the *V* parameter to account for the possibility that microbial abundance influences the relationship between egg size and offspring survival (*i.e.*, the effective *B* parameter) (Fig. S4). As such, the *V* parameter can be used to alter the proportional influence that microbial abundance-a proxy for the functional capacity of that microbial population-has on offspring survival-egg size relationship, with smaller *V* values representing a greater influence on that relationship (Fig. S4).

We introduced several types of microbial manipulators to a population of free-spawning marine invertebrates that had a stable egg size well within range of a typical planktotroph (*i.e.*, 130 µm; Fig. 2)^2,4,8,27^. If the microbial manipulator was only capable of feminization or male killing, then egg size often increased only a few microns (Fig. 2). If the microbe was only capable of enhanced growth, then egg size increased significantly but usually not enough to induce a life-history switch (Fig. 2). If the microbe was capable of manipulation and enhanced growth, then the increase in egg size would most often be sufficient (*i.e.*, >300 µm) to induce a transition from planktotrophy to lecithotrophy (Fig. 2). In this representative example, egg size for these populations became stable after ∼1,700 to ∼2,300 generations, while the distortion in sex ratio caused by the microbial manipulator fixates 46.0x to 53.0x more quickly (Fig. 2, S5). If this microbe is lost following a transition in developmental life-history, then the initial egg size can nearly be recovered and does so 6.5x to 8.0x more quickly than the initial association (Fig. S5).

**Fig. 2:**
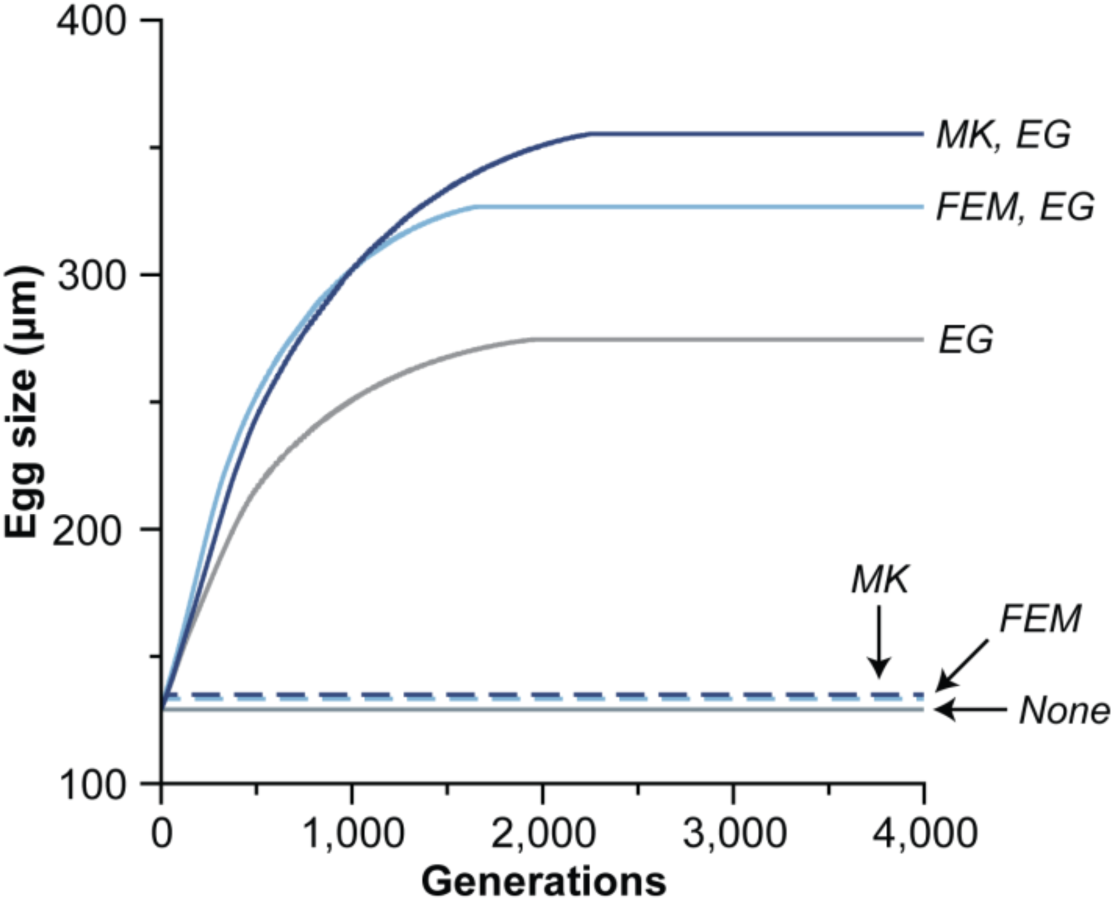
Microbial manipulators enable for transitions in developmental life-history. If a free-spawning marine invertebrate associated with a microbe that has the capacity for feminization (FEM) or male killing (MK), then egg size would often increase a few microns. If this microbe has the capacity for enhanced growth (EG; *i.e.*, a nutritional symbiosis), then egg size usually did not increase enough to induce a transition in developmental life-history. If that microbe has the capacity to manipulate and enhanced growth, then egg size most often would increase enough (*i.e.*, >300 µm^4,27^) that the host undergoes a transition in developmental life-history from the ancestral planktotroph to the derived lecithotroph. This representative example uses the following parameters: *m_k_* = 0.7, *r* = 0.7, *g* = 0.5, *B* =250, and *V* = 280.

### Conditions for microbe-induced transition in developmental life-history

The dynamics described above is an example from a representative set of parameters that demonstrates how a microbial manipulator could induce a transition in developmental life-history in a free-spawning marine invertebrate (Fig. 2). This, however, may not encompass the spectrum of interactions that are observed between marine invertebrates and microbial manipulators. Therefore, we assessed the conditions that microbe-induced transition in developmental life-history are observed. We ran our quantitative genetic population across all possible combinations of *B* values (250 to 750 at increments of 10), *V* values (250 to 750 at increments of 10), and male killing (*m_k_*; 0 to 0.98 at increments of 0.02) or feminization (*r*; 0.5 to 0.99 at increments of 0.01) rates. Initial analyses showed that the presence, not magnitude, of enhanced growth was important and, thus, we only explored no (*g* = 0) or small (*g* = 0.5) enhanced growth (Fig. S6).

Microbe-induced transitions in developmental life-history were observed across a wide range of conditions (Fig. 3, S7). These transitions were more likely to occur when the microbe was capable of manipulation and enhanced growth (57.3% for feminization; 48.7% for male killing), than only enhanced growth or manipulation (21.5% for enhanced growth; 2.2% for only feminization; 1.2% for only male killing) (Fig. 3, S7). Moreover, a more extreme microbe-induced sex ratio distortion is generally needed for a transition in developmental life-history when ecology favors planktotrophy (*i.e.*, as the *B* parameter decreased and the *V* parameter increased) (Fig. 3, S7). Microbe-induced transitions in developmental life-history also led to a wide range of stable egg sizes, of which increased exponentially as a population was manipulated toward female-dominance (Fig. S8-S9). Microbes that are capable of feminization or male killing and enhanced growth increased to an average egg size of 394 μm (± 90 μm, s.d.) and 376 μm (± 76, s.d.) and a maximal egg size of 875 μm and 804 μm, respectively (Fig. 3B, S10). This equates to an average increase in relative egg volume (µm^3^) of 3.9x and 3.1x and maximal increase in relative egg volume of 128.5x and 105.2x, respectively (Fig S10).

**Fig. 3:**
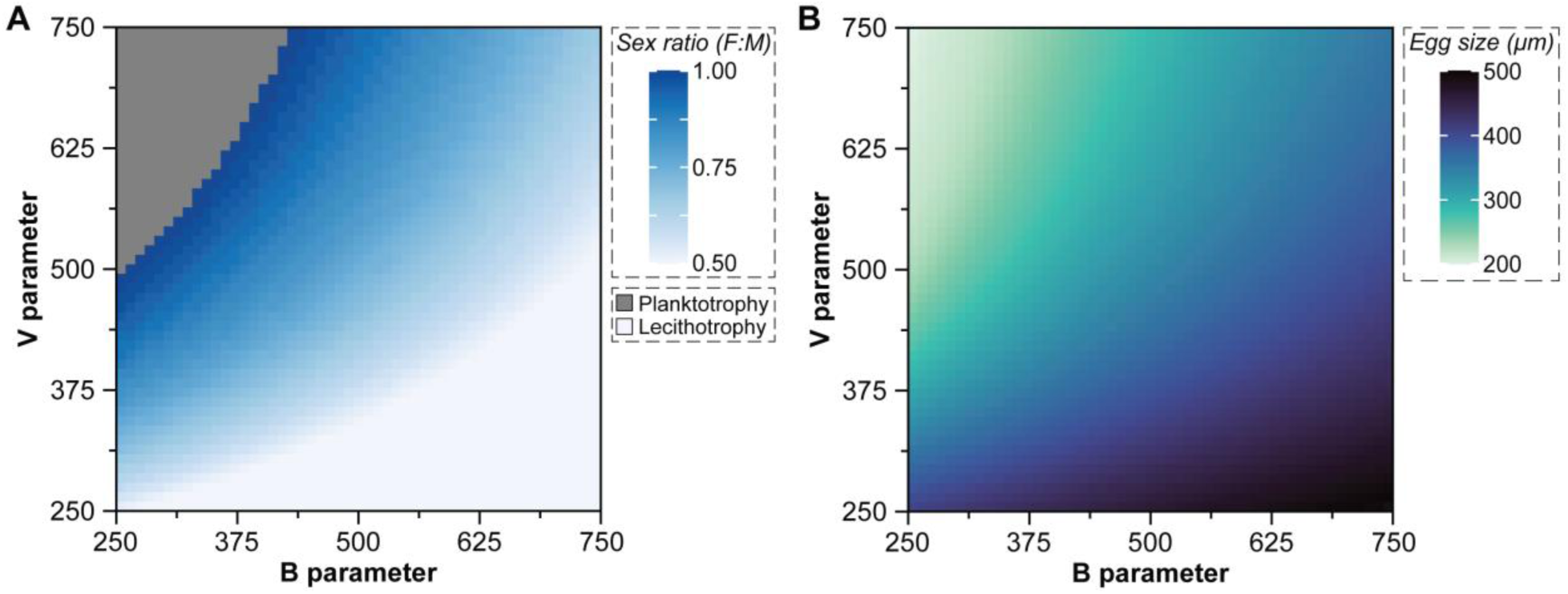
Conditions of a microbe-mediated transition in developmental life-history. The conditions for a microbe-mediate transition in developmental life-history depends on the ecological conditions of the host (*i.e.*, *B* parameter) as well as the manipulating capability of the microbe (*i.e.*, *V* parameter). These factors create a gradient in the sex ratio (A) needed to induce a transition in developmental life-history (*i.e.*, at 300 µm) as well as a gradient in the stable egg sizes that follow this transition (B), with the mean egg size for each combination presented here. No transition may also be observed when ecology heavily favors planktotrophy and the manipulating capability of the microbe is weak (gray, top left of A). When ecology favors larger egg sizes, only enhanced growth is needed for a transition to occur and no manipulation in sex ratio (light blue, bottom right of A). The microbe in this figure represents male killing and enhanced growth, but a nearly identical pattern is also observed for a microbe that has the capacity for feminization and enhanced growth (Fig. S7).

## DISCUSSION

Microbes manipulate host reproduction to enhance the number of microbial cells that are transmitted to the subsequent host generation^17,18,25^. For marine invertebrates, a microbe with the capability to manipulate host reproduction and engage in a nutritional mutualism would associate with a planktotrophic host (Fig. 4). The manipulator would then enhance its fitness by shifting the host’s sex ratio towards female dominance, which would favor larger eggs due to their increased chance of fertilization in a sperm-limited environment (Fig. 4)^31–33^. Egg size of this female-dominant population would then stabilize within the fitness maxima for the host and the microbial manipulator (Fig. 4). A substantial increase in maternal provisioning (*i.e.*, a larger egg) would relax the selective pressures that maintains the feeding structures and accelerate the time to metamorphosis. This, in turn, should lead to a derived-likely lecithotrophic-developmental life-history^7,8,34,35^. Shortening this developmental window would limit the genetic exchange between populations, reduce the geographic range, and increase speciation and extinction rates^4,8,9,27,36,37^. Microbial manipulators now represent a plausible mechanism that can drive a switch between the major developmental life-history amongst marine invertebrates.

**Fig. 4:**
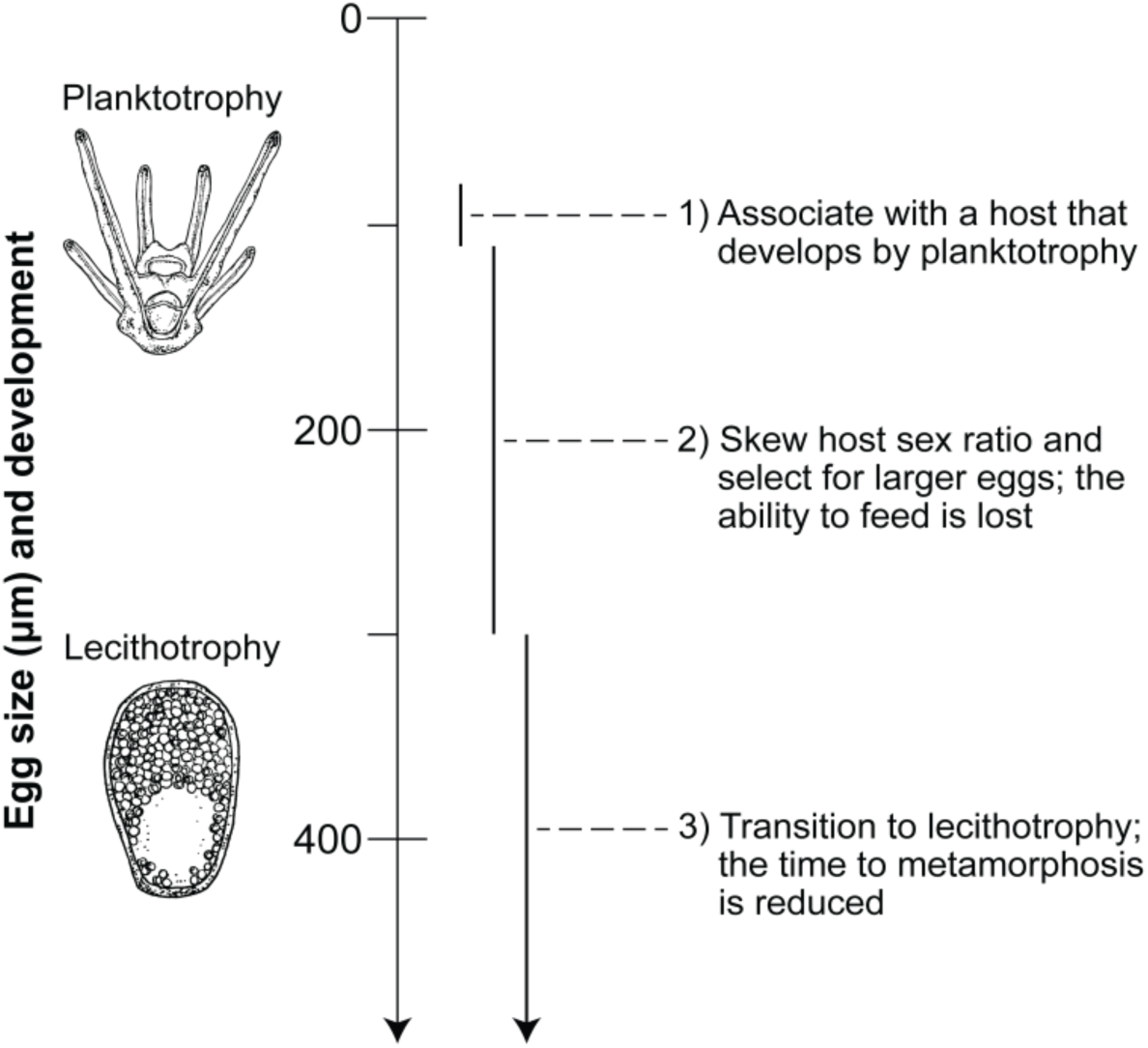
Microbes as manipulators of developmental life-history. A microbial manipulator associated with a planktotrophic (feeding) host would enhance its fitness by shifting the sex ratio towards female dominance. This favors larger eggs due to the increased chance of fertilization in a sperm-limited environment. Egg size in this female-dominant population would then stabilize between the fitness maxima of the host and the microbial manipulator. A substantial increase in maternal investment would relax the selective pressures that maintain the feeding structures and accelerate the time to metamorphosis. This, in turn, would lead to a derived-likely lecithotrophic (non-feeding)-developmental life-history. Processes relating to the host reflects previous demonstrations of how development changes during life-history transitions^4,7,49^. Drawings are by Alexandra Hahn.

Egg size, developmental mode, phylogeny, and geography can be used to identify potential microbial manipulations of developmental life-history in marine invertebrates^7,12,27,38–40^. We identified candidates from 17 genera representing annelids, echinoderms, and molluscs and from most major oceans (Table 1, S4). These were predominately echinoderms (82.4%; *e.g.*, the sea star *Parvulastra*^38^, the sea urchin *Heliocidaris*^20^, and the brittle star *Macrophiothrix*^41^) and the majority were from Oceania (58.8%) (Table 1, S4), an area of the world’s oceans with the richest diversity of developmental life-histories^12,38^. More than 80% of marine invertebrates are predicted to associate with microbial manipulators^22^ and, thus, available natural history data and well-resolved, species-level phylogenies for many marine invertebrate phyla likely limit our capacity to identify additional candidates. Therefore, we hypothesize that microbial manipulators have a yet-to-be appreciated influence on the reproductive strategies and developmental life-history of marine invertebrates.

**Table 1:**
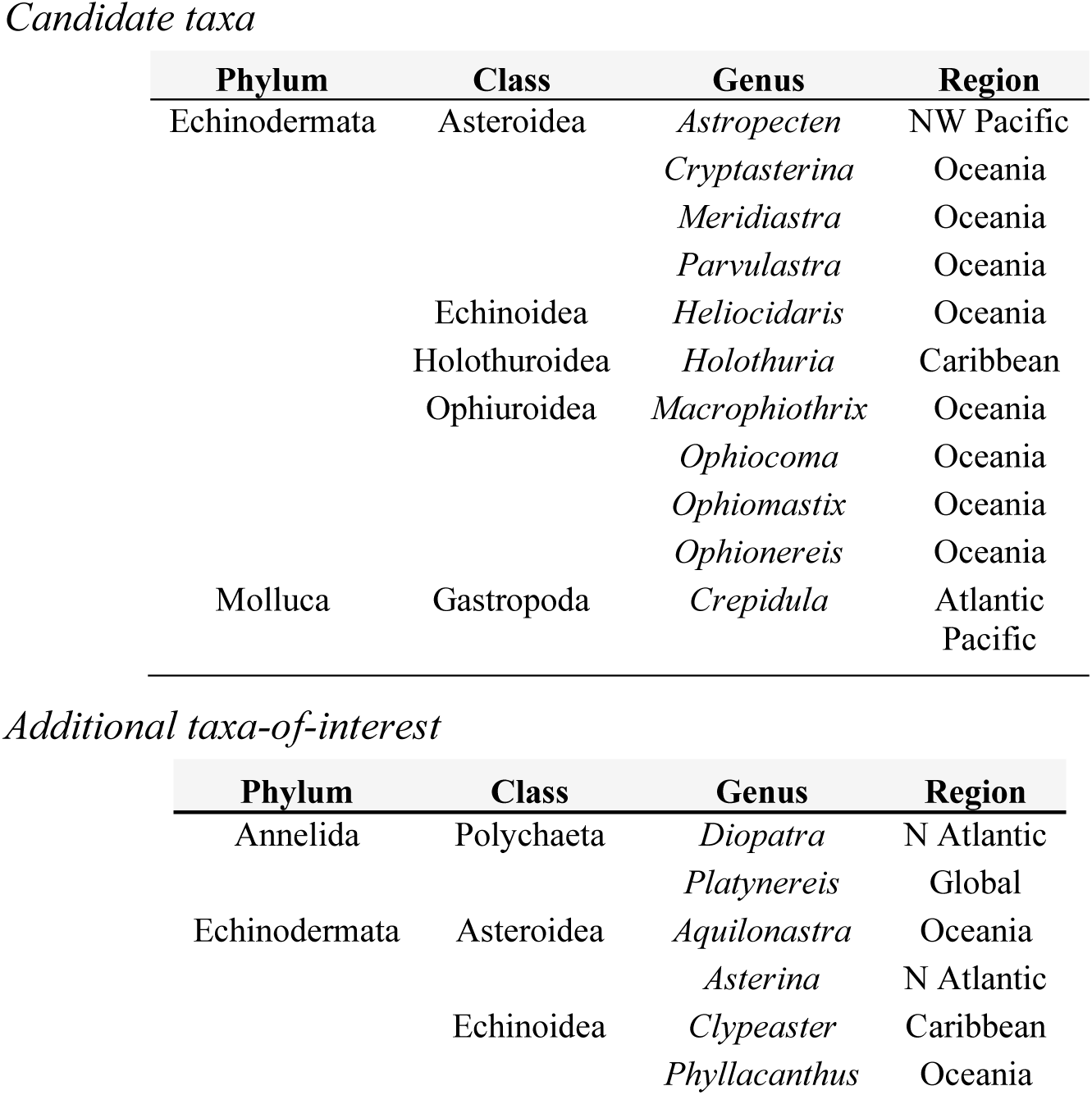
Candidate marine invertebrate genera of a microbe-mediate transition in developmental life-history (see, Table S4 for references).

We describe a set of circumstances under which marine invertebrates would transition between developmental life-histories, but the limitations of this model must also be considered. First, the biological data used to parameterize this models come primary from sea urchins because of their widespread use in embryology, evolution, and ecology^22,42,43^. Therefore, the implication of this model are more bias towards sea urchins than other groups of free-spawning marine invertebrates (*e.g.*, annelids or gastropods)^1,6,8^. Second, transitions in developmental life-history were modeled based on an egg size of 300 μm. This is, thus, an indirect assessment of the complexities involved in developmental evolution that is most strictly adhered to by sea urchins^4,27^. The specific conditions for microbe-induced transition in developmental life-history would be expected to differ between major groups of marine invertebrates. Third, we only allowed for egg size to evolve in sperm limiting conditions. What this model does not consider is how sperm would also adapt (*e.g.*, affinity, chemotaxis, longevity, or quantity) to enhance fertilization in these conditions^31,32^.

If microbes act as an evolutionary agent responsible for switches in the reproductive strategy of marine invertebrates, then it remains paramount to understand two fundamental questions. First, are there transitions in developmental life-history that are and are not the result of a microbial manipulator and, if so, how do they differ? Second, are the mechanisms of manipulation by terrestrial and marine microbes convergent or divergent? In the former, our candidate genera are far below reported estimates for transitions in developmental life-history amongst marine invertebrates^1,6,7,27,38^. In the latter, *Echinorickettsia raffii*-a close relative of *Wolbachia*-is hypothesized to use an ankyrin protein that has homology with the male killing gene of terrestrial microbes to have manipulated the sea urchin *Heliocidaris*^20,44^. It would appear that transitions in developmental life-history could occur with and without reproductive manipulators and that there may be some genomic convergence between the microbial relatives in these environments. We anticipate that acknowledging microbial manipulators as a factor underlying the ecological and evolutionary processes governing marine invertebrate reproduction, development, and life-history will shed light on age-old questions and conundrums in similar ways that they have for terrestrial life^2–4,8,16–18^.

## METHODS

### Fertilization dynamics

We used a modified version of the Styan^43^ polyspermy kinetic model to determine the fertilization dynamics of free-spawning marine invertebrates^22^. Specifically, we converted egg (*E_T_;* egg per μL) and sperm (*S_T_*; sperm per μL) concentrations to be explicitly determined by female and male density. We then determined the average number of sperm that may potentially fertilize an egg during a free-spawning event, *x*. This is a function of fertilization efficiency (*F_e_*), sperm concentration (*S_T_*; sperm per μL), egg concentration (*E_T_;* egg per μL), the biomolecular collision constant (β_0_; mm^3^ per s), and the sperm half-life (τ; s):

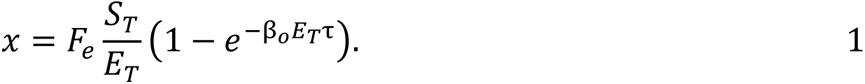

Moreover, the biomolecular collision constant (β_0_) was determined by the cross-sectional area of the egg (σ; mm^2^) and sperm velocity (*v*; mm per sec):

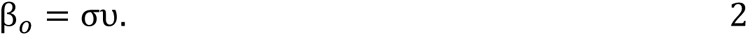

The biomolecular collision constant and the time to elicit a block to polyspermy (*t_b_*; s) was then used to calculate the average number of extra sperm that may contact a fertilized egg, which then results in polyspermy (*b*):

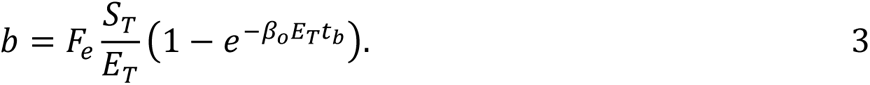

The proportion of eggs that were fertilized by a single sperm and successfully develop was estimated by:

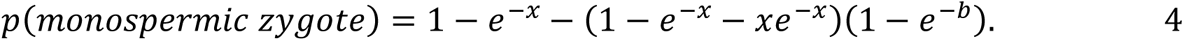

Zygote density (*N_z,t_*; zygotes per μL) was then be estimated based on the probability of a monospermic zygote (Equation 4) and total egg density (*E_T_*):

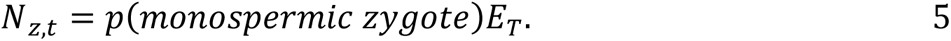

A summary of the variables and parameters are provided in Table S1.

### Population dynamics

We integrated the modified polyspermy kinetic model into a population model to simulate how a microbe that manipulates reproduction by feminization or male killing and can compensate host fitness by supplementing host nutrition affects population size^22^. This was a discrete time population model with overlapping generations. Reproduction, development, and adult mortality are assumed to occur in that order because marine invertebrates tend to spawn seasonally^45^.

Offspring survival based on egg size (*s*_σ_) was determined using the Vance-Levitan survival function^3,26,42^, where survival was determined by the survival function parameter (*B*) and egg size (a):

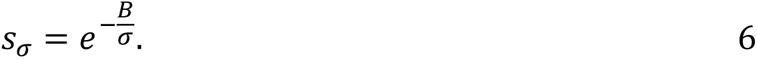

Adult mortality (*M_a_*) was assumed to be density-dependent and based on the maximum adult mortality rate (*m_a_*), carrying capacity (*k*; individuals per m^2^), total density of the population at time *t* (*N_t_*; individuals per m^2^), and the degree to which density influences mortality (*d*; *i.e.*, the shape parameter). Both the current adult density and density of surviving zygotes that settled were assumed to influence density dependence, where *N^’^_z,t_* is the density of zygotes (zygotes per μL) that survived after a microbe-induced male killing, and ψ (μL per m^2^) is the settlement constant that describes the proportion of surviving zygotes that transitioned from larvae to juveniles:

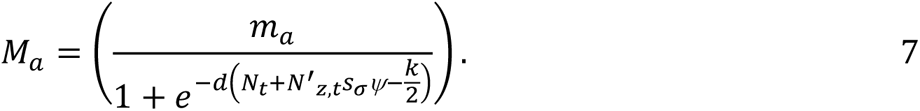

The density of females in the next generation (*N_f,t+1_*; individuals per m^2^) was then determined by the density of surviving female zygotes (*N_z,f,t_*; zygotes per μL) that settled plus the density of surviving female adults at generation *t* (*N_f,t_*; individuals per m^2^):

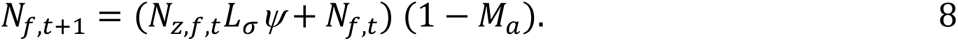

For male killing scenarios, we assumed that females produced an equal number of daughters and sons and, therefore, the density of female zygotes (*N_z,f,t_*) is calculated by dividing the density of zygotes (*N_z,t_*) by two. The density of males in the next generation (*N_m,t+1_*; individuals per m^2^) has the same structure as Equation 8. The density of male zygotes (*N_z,i,m,t_*; zygotes per μL) is influenced by mortality due to male killing (*m_k_*):

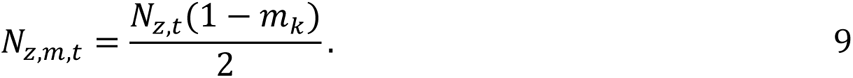

For feminization scenarios, sex ratio of zygotes were biased by feminization rate *r*. Such that the density of female zygotes (*N_z,f,t_*) was:

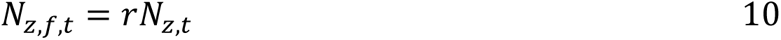

and the density of male zygotes (*N_z,i,m,t_*) was:

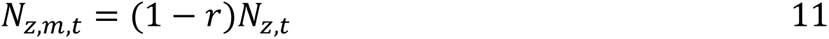

A summary of the variables and parameters are provided in Table S2.

### Invasion grid model

We performed an invasion grid analysis to simulate the population dynamics for free-spawning marine invertebrates along a continuum of egg sizes (*σ*)-from 50 µm to 1,500 µm at increments of 10 µm^12^-until an equilibrium was reached (*i.e.*, a change in population densities was equivalent to zero) for a population with or without a microbe that can manipulate reproduction by feminization or male killing and/or enhanced growth. We then calculated the number of surviving fertilized eggs (*i.e.*, fitness) of an individual that produced the resident egg size-the current position on the grid-and compared this to the fitness of a mutant individual that produced an egg size that was either 10 µm smaller or larger. Fitness of a given egg size (*W_σ_*) was a function of the probability of fertilization success (*P_f,σ_*), egg density (*E_σ_*; egg per µL), and survival (*L_σ_*) (Fig. 1A):

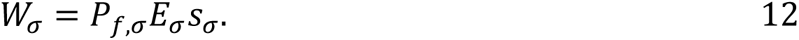

The probability of fertilization success was calculated using the resident egg number because we assume that the mutant is rare^26^, but egg size was varied based on whether the calculation was for a resident, smaller mutant, or larger mutant. Egg size is inversely proportional to fecundity^46^ and, thus, we calculated the density of eggs produced using a constant for planktotrophs (*i.e.*, 1.13^22^) divided by egg size (*σ*) and multiplying that by the enhanced growth rate factor (*g*). Specifically, that *B* is the parameter determining the importance of egg size on survival, where larger values of *B* equates to egg size being is more important to offspring survival (Fig. S1A). We performed this invasion grid analysis to solve for the optimal egg size across male killing rates (0 to 0.99 at increments of 0.03), feminization (0.5 to 0.995 at increments of 0.015), enhanced growth rates (0 to 3 at increments of 0.1), and *B* values (100 to 3,000 at increments of 100).

We then calculated fitness for the host across all model parameters to determine this fitness landscape. Host fitness was calculated using Equation 12 assuming the evolutionary stable egg size. Fitness values were then compared between a host that did (*W_i,_σ*) and did not (*W_u,_σ*) associated with a microbial manipulator to assess how an evolutionary shift in egg size changed the fitness landscape:

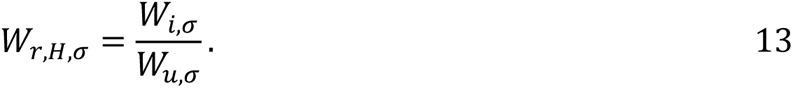

### Simulation of egg size evolution

We developed a quantitative genetic population model to numerically simulate the evolution of egg size (*σ*) in a population of free-spawning marine invertebrates. To approximate a continuous normal distribution of female egg size, we created bins of egg sizes (*σ*) that were 0.2 µm in size, which resulted in 14,950 bins across the range of possible egg sizes in the model (10 to 3,000 µm). We initialized populations with a mean of 180 µm, a standard deviation of 9, and a range of ±10 standard deviations from the mean. To generate starting numbers of individuals at each egg size bin, we calculated the relative Gaussian probability density function multiplied by the total population size (1 individual per m^2^). Males in the population were assumed to not vary in any reproductive trait.

Fitness for each egg size bin was a product of fertilization success (Equation 5), larval survival based on egg size (Equation 6), and the settlement constant ψ (μL per m^2^). To account for a potential for microbial abundance to influence the relationship between egg size and offspring survival, we additionally modeled scenarios of enhanced growth (*g* > 0), where egg size influenced the effective *B* parameter (*B_v_*) because larger eggs would have higher microbial abundance^28–30^. We modeled this relationship with a Michaelis-Menten equation, such that the *V* parameter represents the egg size where the *B* parameter is influenced halfway to maximal possible influence (Fig. S4A)^47^:

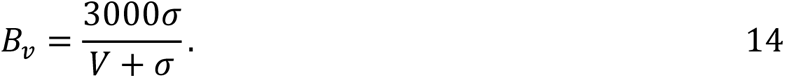

In these scenarios where microbial abundance influences the effective *B* parameter (*B_v_*), larval survival (Equation 6) is modified by adding *B_v_* and *B* (Fig. S4B):

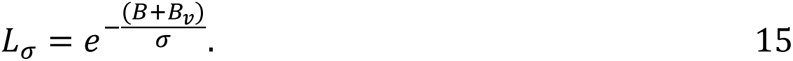

After fitness (*i.e.*, the number of surviving offspring) was calculated for each egg size bin, we calculated the egg size distribution of offspring using standard quantitative genetics^48^. Specifically, the selection differential on egg size (*S_σ_)* was calculated as a function of mean fitness 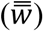 and covariation of fitness and egg size (*σ*):

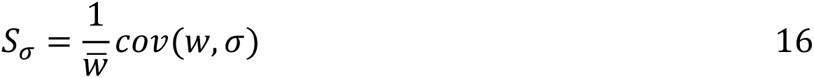

The response to selection (*R*) was calculated as the heritability (*h^2^*) multiplied by the selection differential (*i.e.*, the quantitative genetic breeder’s equation):

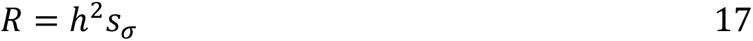

We assumed that heritability was constant at 0.05. Preliminary analyses varying heritability influenced the quickness of egg size evolution and not stable egg size outcomes. We then generated the female offspring distribution with a mean of the previous generation + *R*, a standard deviation of 9, and a total density equal to the density of surviving female offspring produced accounting for either feminization or male killing and enhanced growth.

For male killing scenarios, we assumed half of surviving offspring were males and the other half females. The surviving males experienced a male killing mortality rate (*m_k_*). For feminizing scenarios, where *r* represents the feminization rate: *r* proportion of surviving offspring were female and 1-*r* surviving offspring were male. The density of each egg size bin for females in the next generation was added to surviving individuals from the previous generation. The density of males in the next generation was calculated by adding the density of surviving male offspring to density of survival males from the previous generation. For simplicity, we assumed a constant adult mortality rate (*m_a_*) of 0.9. Initial simulations varying this number did not qualitatively affect the results. If total population size after mortality exceeded the set carrying capacity (*k*), population size was adjusted to *k*. Simulations were run for 4,000 generations as mean egg size stabilized well before then across a wide range of parameters. Fertilization parameters were the same as the invasion grid model (Table S1). Summary of parameter values specific to the quantitative genetic model are given in Table S3.

### Identification of candidate host taxa

We used the literature on marine invertebrate egg size^12,27,40^, developmental mode ^12,27,40^, phylogeny^38,39^, and geographic distribution (via World Register of Marine Species) to identify candidate transitions in developmental life-history that could have been the result of a microbial manipulation. We define a candidate microbial manipulation of developmental life-history as the ancestral host species having an egg size of less than 225 µm [*i.e.*, an approximate threshold for obligate planktotrophy^4,27^ and near the highest egg size within our minimal relative fitness (Fig. 1C)] and the derived species having an egg size between 300 and 500 µm [*i.e.*, the smallest egg size for lecithotrophy and within the peak in relative fitness (Fig. 1C)]. Moreover, sister species also have to inhabit a similar geographical region and exhibit a transition in developmental mode (see, Table 1, S4). Additional taxa-of-interest were genera that nearly met these qualifications (see, Table 1, S4).

## REPORTING SUMMARY

Further information on research design is available in the Nature Research Reporting Summary linked to this article.

## DATA AVAILABILITY

All data to support the analyses and conclusions of this study are available online.

## CODE AVAILABILITY

All computer code to support the analyses and conclusions of this study are available online.

## ACKNOWLEDGEMENTS

We thank Jussi Lehtonen (Univ. Jyväskylä) for insightful conversations that led to the completion of this model as well as Mary Sewell (Univ. Auckland) and Craig Young (Univ. Oregon) for providing access to their datasets on egg size in echinoderms. We are grateful for Alexandra Hahn (GEOMAR Helmholtz Centre for Ocean Research) for drawing the sea urchin larvae used in Fig. 4. M.C.K. was supported by an NSF Graduate Research Fellowship, and T.J.C. was supported by a postdoctoral fellowship from the Alexander von Humboldt Foundation, the German Research Foundation (project 261376515; CRC 1182, “Origin and Function of Metaorganisms”), and the GEOMAR Helmholtz Centre for Ocean Research.

## AUTHOR CONTRIBUTIONS

M.C.K. and T.J.C. conceived and designed the study. M.C.K. performed the modeling. M.C.K. and T.J.C. interpreted the model. T.J.C. assessed natural history data. M.C.K. and T.J.C. wrote and edited the manuscript.

## COMPETING INTERESTS

The authors declare no competing interests.

## Supplementary Information

### Supplementary Tables

**Table S1.**
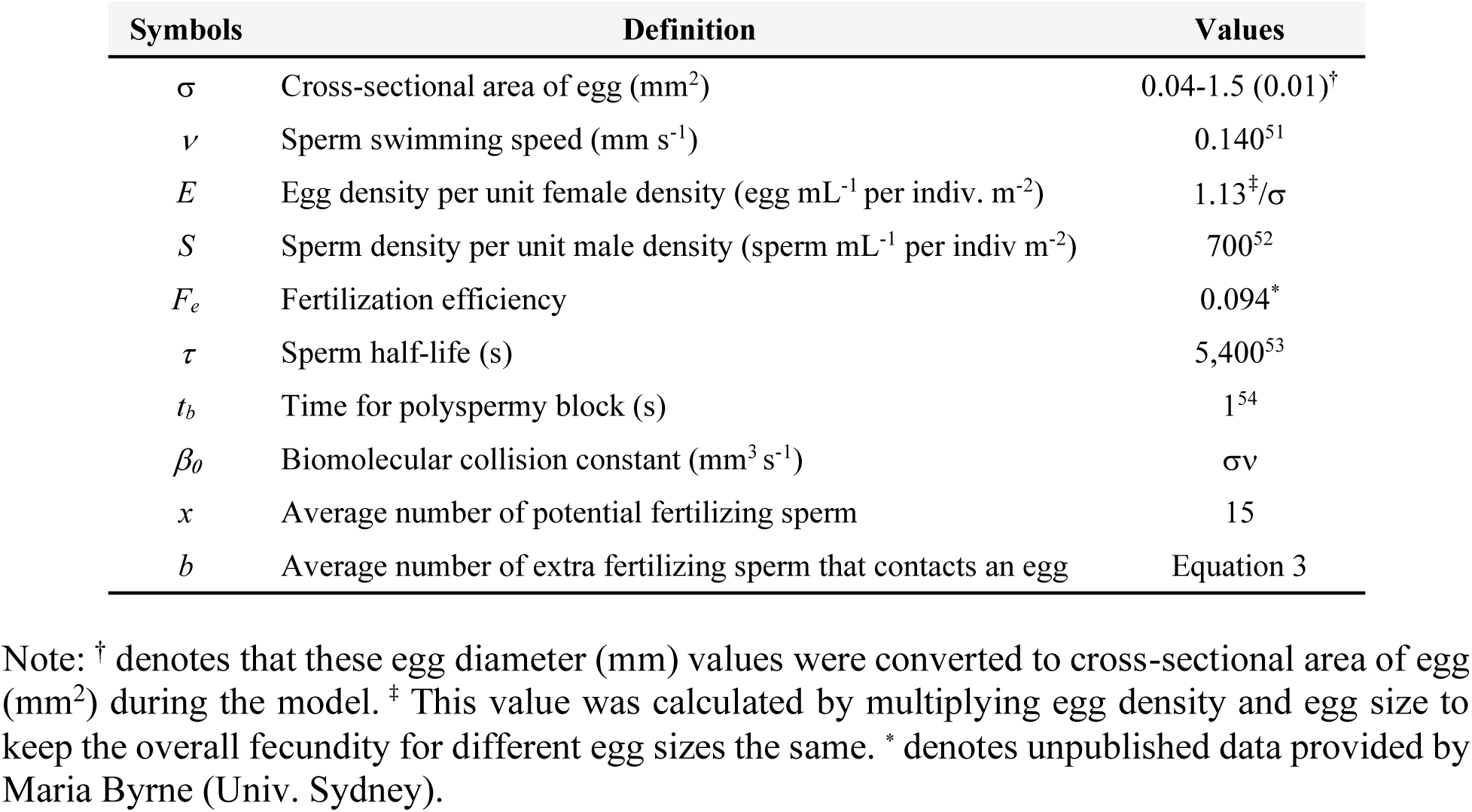
Definitions and values of parameters for fertilization dynamics adapted from Styan^50^.

| Symbols | Definition | Values |
| --- | --- | --- |
| $\sigma$ | Cross-sectional area of egg (mm<sup>2</sup>) | 0.04-1.5 (0.01)<sup>†</sup> |
| $v$ | Sperm swimming speed (mm s<sup>-1</sup>) | 0.140<sup>51</sup> |
| $E$ | Egg density per unit female density (egg mL<sup>-1</sup> per indiv. m<sup>-2</sup>) | 1.13<sup>‡</sup>/$\sigma$ |
| $S$ | Sperm density per unit male density (sperm mL<sup>-1</sup> per indiv m<sup>-2</sup>) | 700<sup>52</sup> |
| $F_e$ | Fertilization efficiency | 0.094<sup>*</sup> |
| $\tau$ | Sperm half-life (s) | 5,400<sup>53</sup> |
| $t_b$ | Time for polyspermy block (s) | 1<sup>54</sup> |
| $\beta_0$ | Biomolecular collision constant (mm<sup>3</sup> s<sup>-1</sup>) | $\sigma v$ |
| $x$ | Average number of potential fertilizing sperm | 15 |
| $b$ | Average number of extra fertilizing sperm that contacts an egg | Equation 3 |

Note: <sup>†</sup> denotes that these egg diameter (mm) values were converted to cross-sectional area of egg (mm<sup>2</sup>) during the model. <sup>‡</sup> This value was calculated by multiplying egg density and egg size to keep the overall fecundity for different egg sizes the same. <sup>\*</sup> denotes unpublished data provided by Maria Byrne (Univ. Sydney).

**Table S2.** Definitions and values of parameters for non-stage-based population dynamics model.

| Symbols | Definition | Values |
| --- | --- | --- |
| Parameters |  |  |
| $k$ | Adult carrying capacity (indiv. m<sup>-2</sup>) | 100 |
| $\psi$ | Settlement constant (mL m<sup>-2</sup>) | 1 |
| $d$ | Mortality rate of change | 0.1 |
| $m_a$ | Max adult mortality | 0.99 |
| $B$ | Importance of egg size | 300 – 3000 (300) |
| $g$ | Enhanced growth (fecundity enhancement, %) | 0-3 (0.1) |
| $r$ | Feminization rate | 0 – 0.99 (0.03) |
| $m_k$ | Male killing rate | 0.5 – 0.99 (0.06) |
| Functions |  |  |
| $L\sigma$ | Larval survival | Equation 6 |
| $M_a$ | Adult mortality | Equation 7 |
| Variables |  |  |
| $N_t$ | Density of all adult indiv. at time t | 0.2 |
| $N_{f,t}$ | Density of adult females at time t | 0.1 |
| $N_{m,t}$ | Density of adult males at time t | 0.1 |
| $N_{z,t}$ | Density of initial zygotes at time t | N/A |
| $N_{z,t}^*$ | Density of zygotes at time t after male killing | N/A |
| $N_{z,f,t}$ | Density of new females at time t | N/A |
| $N_{z,m,t}$ | Density of new males at time t | N/A |

Note: Capital letters are functions or variables, and initial values are given when available. Zygote density units are zygotes per $\mu\text{L}$ and adult density units are individuals per $\text{m}^2$ . Numbers in parenthesis indicate step size in sensitivity analyses.

**Table S3.** Definitions and values of parameters for quantitative genetic population model.

| Symbols | Definition | Values |
| --- | --- | --- |
| Parameters |  |  |
| $k$ | Adult carrying capacity (indiv. m<sup>-2</sup>) | 2.6 |
| $\psi$ | Settlement constant (mL m<sup>-2</sup>) | 100 |
| $m_a$ | Adult mortality | 0.9 |
| $h^2$ | Heritability | 0.05 |
| $B$ | Importance of egg size | 250-750 (10) |
| $g$ | Enhanced growth (fecundity enhancement, %) | 0 – 2 (0.5) |
| $m_k$ | Male killing rate | 0-0.98 (0.01) |
| $r$ | Feminization rate | 0.5-0.99 (0.01) |
| $V$ | Rate parameter determining how egg size influences $B$. | 250-750 (10) |
| Functions |  |  |
| $B_v$ | Effective $B$ parameter modifier | Equation 14 |
| $L_\sigma$ | Larval survival | Equation 15 |
| $S_\sigma$ | Selection differential | Equation 16 |
| $R$ | Response to selection | Equation 17 |

Note: Capital letters are functions or variables, and initial values are given when available. Zygote density units are zygotes per $\mu\text{L}$ and adult density units are individuals per $\text{m}^2$ . Numbers in parenthesis indicate step size in sensitivity analyses.

**Table S4.**
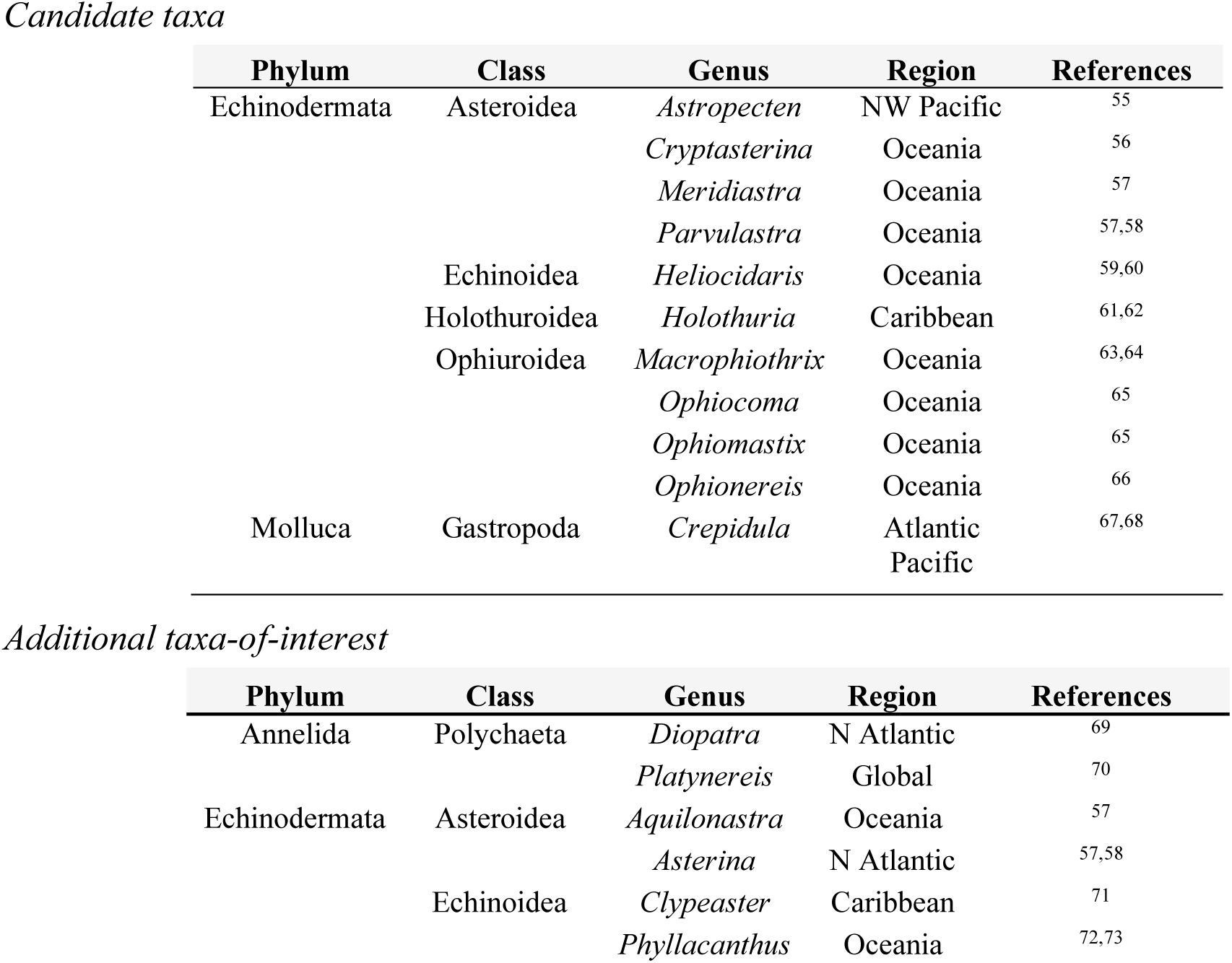
References for candidate marine invertebrate genera of a microbe-mediated transition in developmental life-history.

### Supplementary Figures

**Fig. S1:**
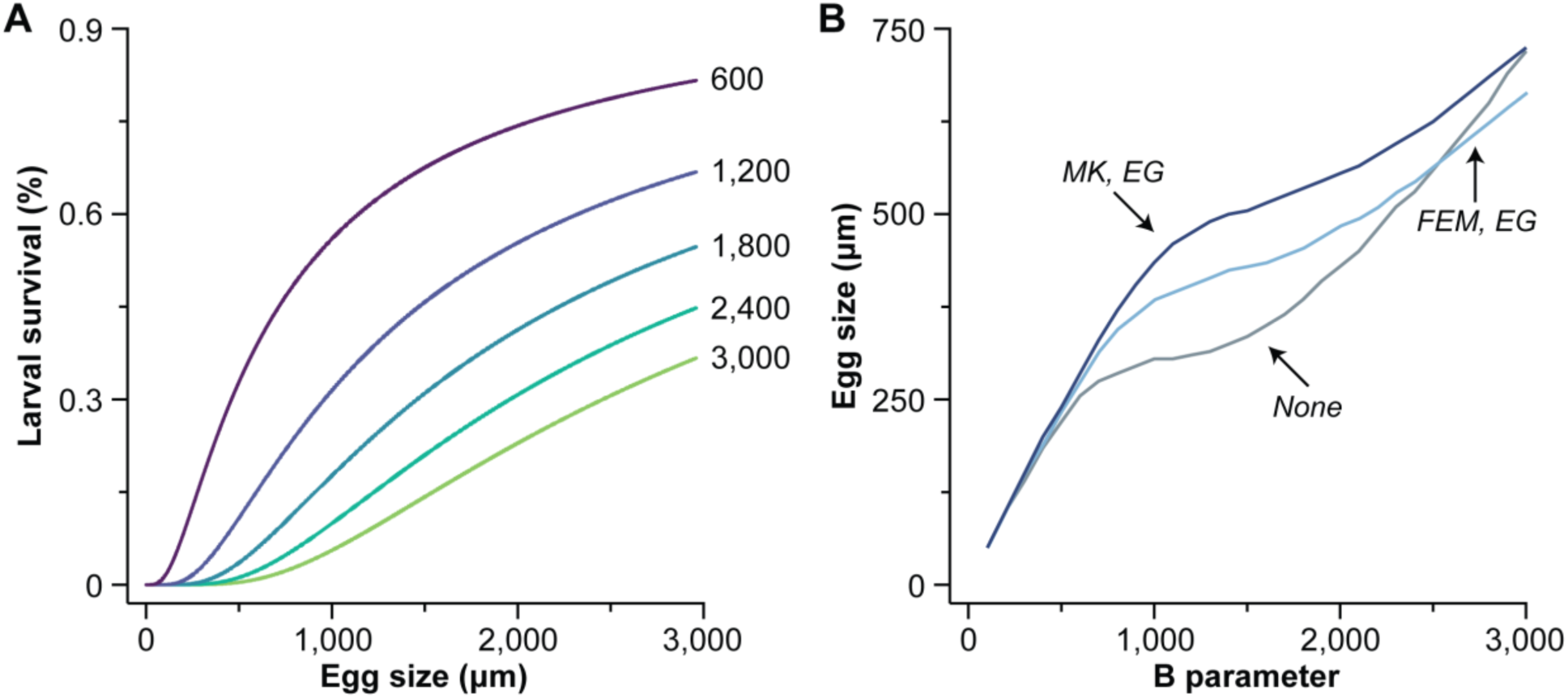
Relative importance of *B* parameter to egg size and larval survival. (A) The relationship between egg size and larval survival at *B* values from 600 to 3,000 using Equation 6. (B) Stable egg size (µm in diameter) across *B* parameter (*i.e.*, the offspring survival function) values for populations without a manipulator (gray) compared to those with a feminizing (light blue) or male killing (dark blue) manipulator that also enhances growth.

**Fig. S2:**
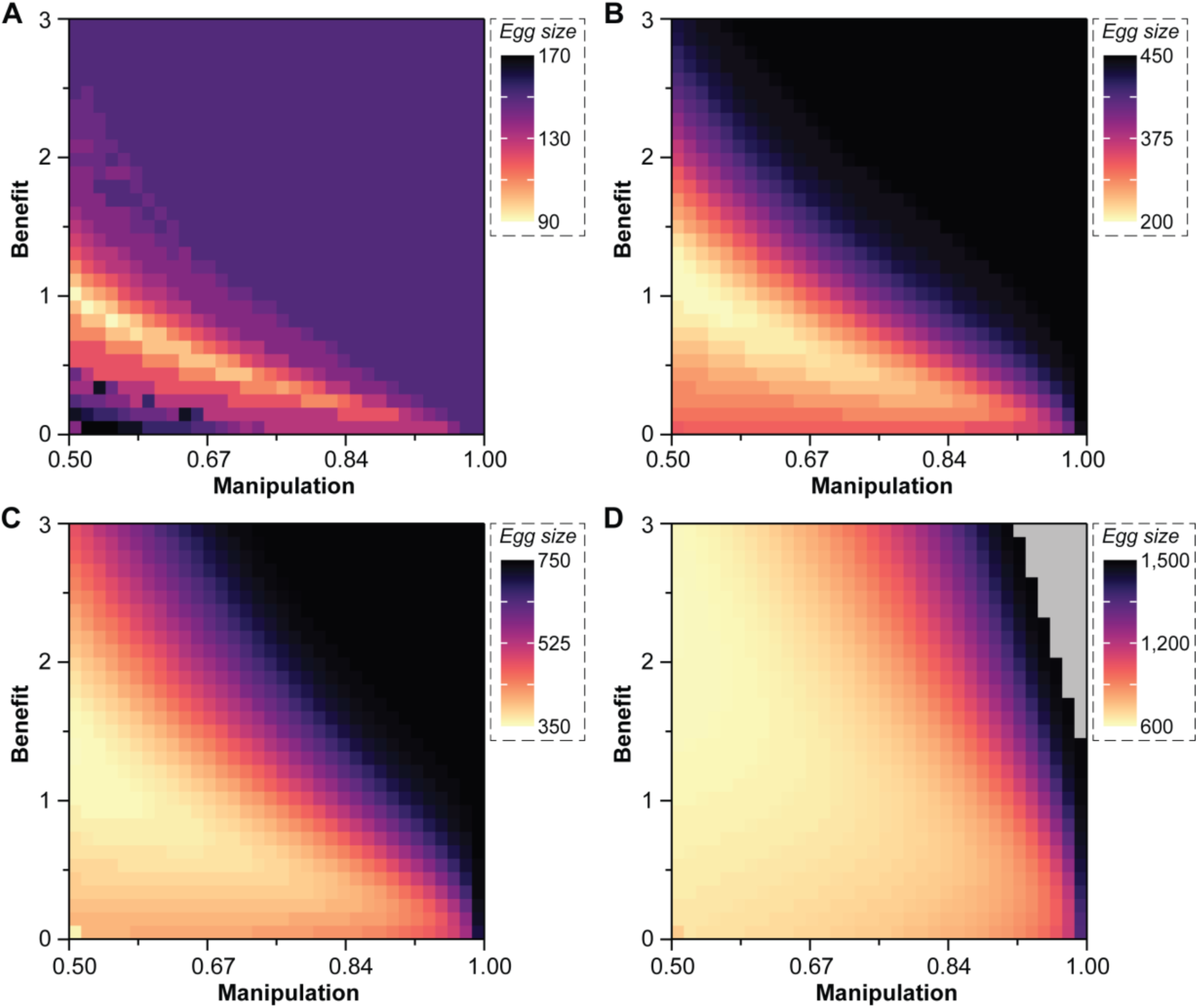
Influence of *B* parameter, feminization, and enhanced growth on egg size evolution. Evolutionary stable egg size (µm in diameter) across feminization (0.50 to 0.99 at increments of 0.03) and enhanced growth rates (0 to 3.0 at increments of 0.1) when the *B* parameter equals 300 (A), 900 (B), 1,500 (C), or 3,000 (D). Evolutionary stable egg sizes were determined using the invasion grid analysis model. Grey shading indicates parameter combinations that resulted in unviable populations.

**Fig. S3:**
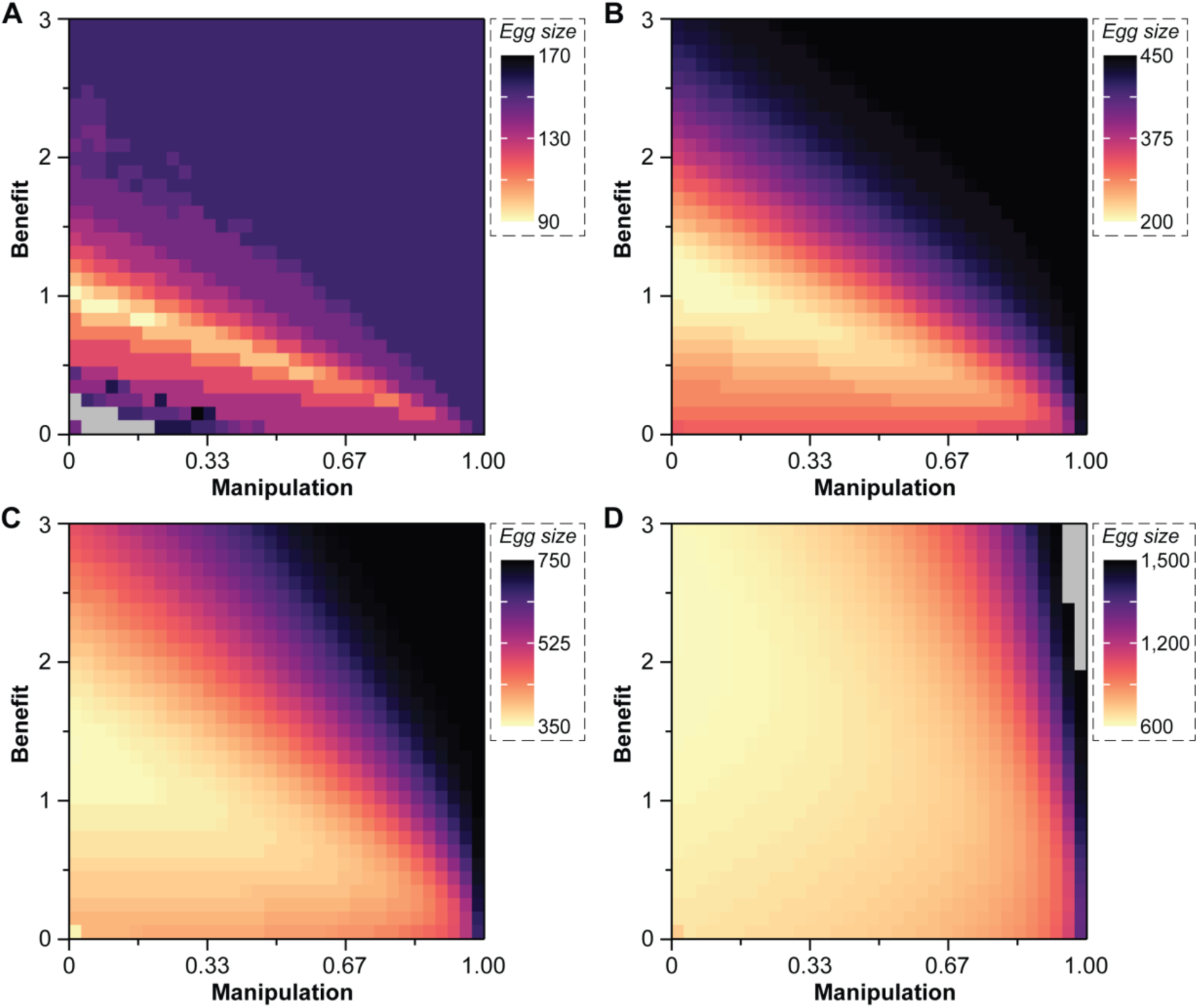
Influence of *B* parameter, male killing, and enhanced growth on egg size evolution. Evolutionary stable egg size (µm in diameter) across male killing (0 to 0.99 at increments of 0.03) and enhanced growth rates (0 to 3.0 at increments of 0.1) when the *B* parameter equals 300 (A), 900 (B), 1,500 (C), or 3,000 (D). Evolutionary stable egg sizes were determined using the invasion grid analysis model. Grey shading indicates parameter combinations that resulted in unviable populations.

**Fig. S4:**
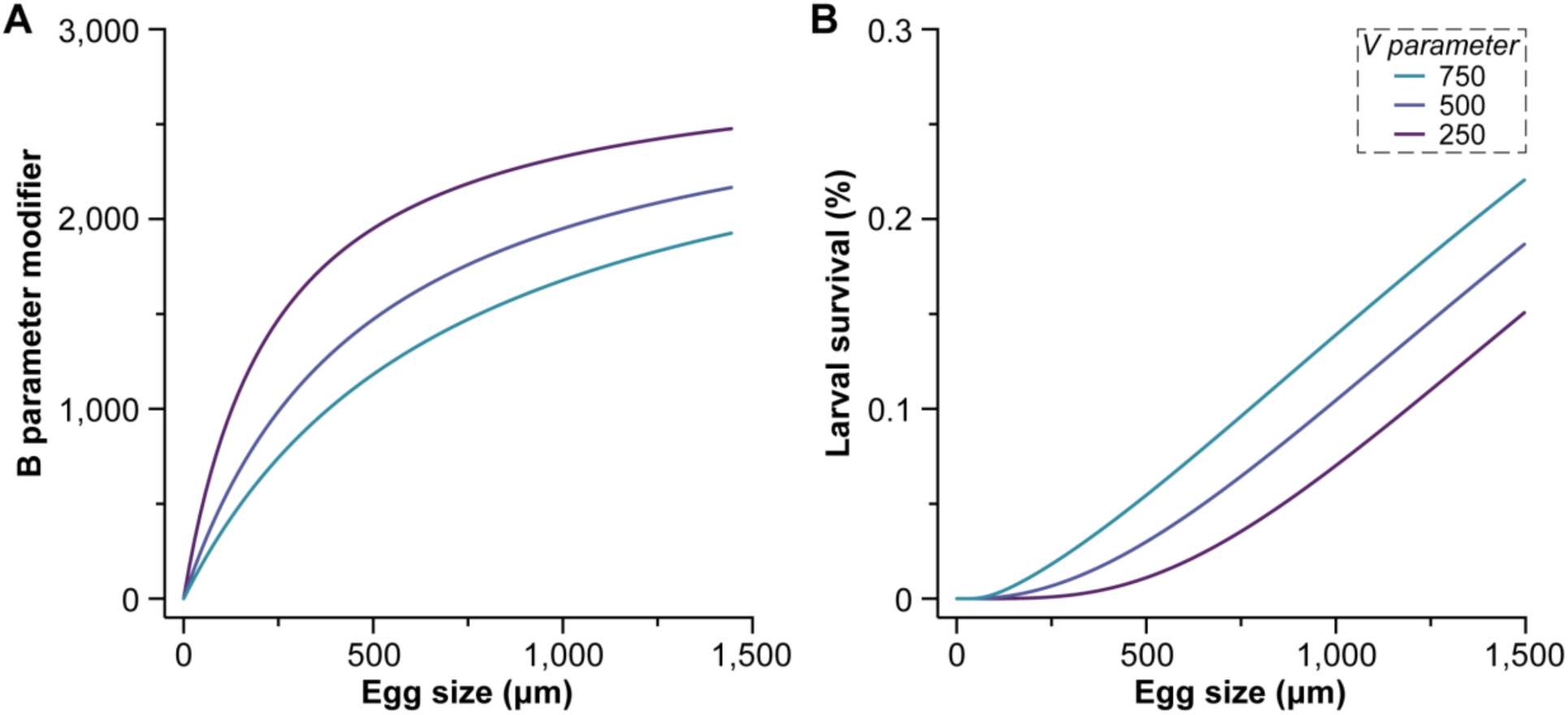
*V* parameter determines how egg size modifies the *B* parameter and the relationship between egg size and larval survival. (A) The relationship between egg size and how the *B* parameter is modified at different *V* values using Equation 14. (B) The relationship between egg size and larval survival at different *V* values when *B* = 250 (Equation 15).

**Fig. S5:**
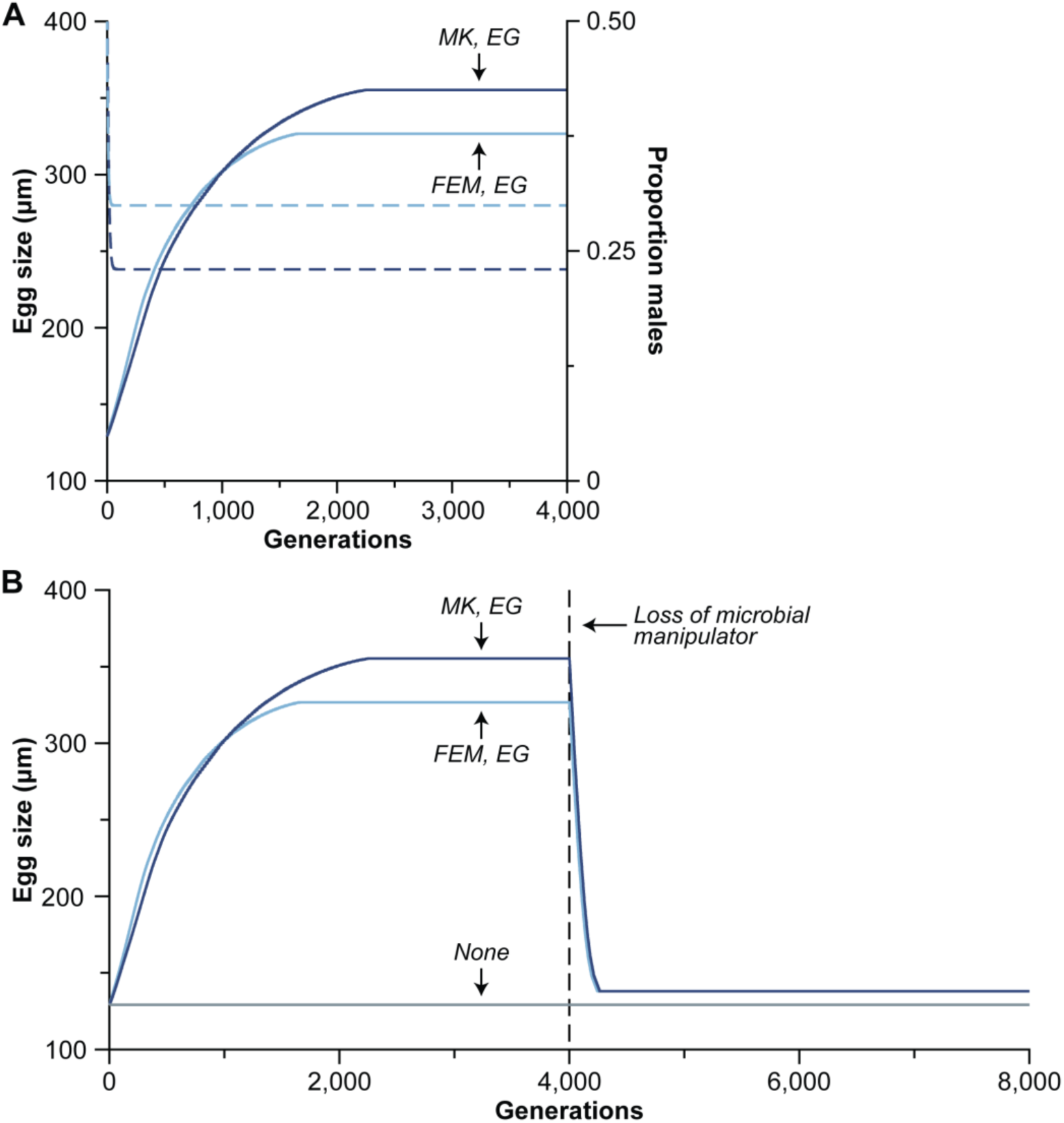
Speed and reversal of a microbe-induced transition in developmental life-history. (A) If a free-spawning marine invertebrate associates with a microbe that has the capacity for male killing (MK; dark blue; solid line) or feminization (FEM; light blue; solid line) and has the capacity for enhanced growth (EG; *i.e.*, a nutritional symbiosis), then egg size (µm in diameter) increases enough that the host can undergo a transition in developmental life-history. This change in egg size and developmental life-history is preceded by a shift in sex ratio (dashed lines) that is caused by the microbial manipulator. (B) Removal of a microbe that manipulates by feminization or male killing and enhanced growth from a stable population following the transition in developmental life-history. Egg size following removal of the microbial manipulator nearly results in a return to a population that never associated with a microbial manipulator (gray). Parameter values are the same as Fig. 2 before loss of microbial manipulator.

**Fig. S6:**
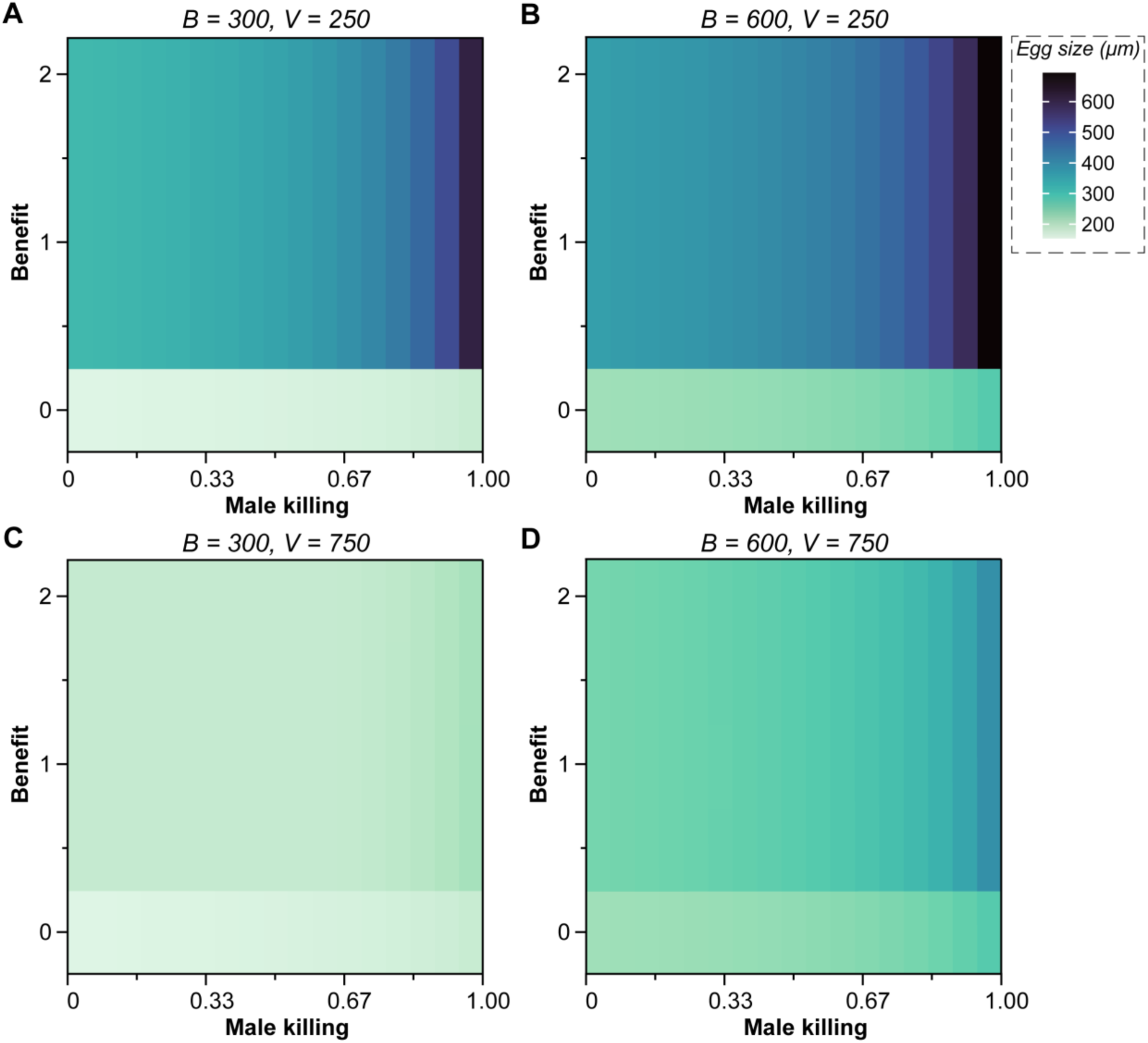
Presence, not magnitude, of enhanced growth does not influence stable egg size. Heat map of stable egg sizes at different male killing rates and enhanced growth factors across different combinations of *B* and *V* values.

**Fig. S7:**
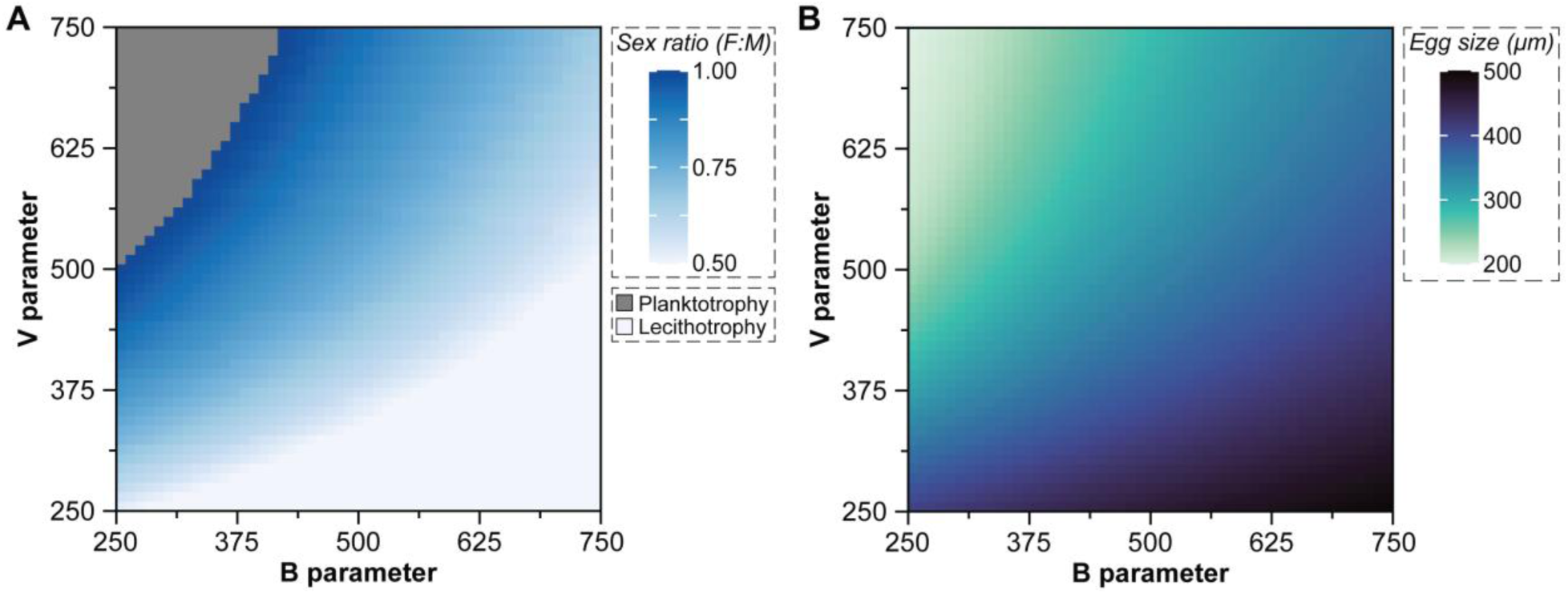
Feminization and enhanced growth results in similar patterns as male killing and enhanced growth. The conditions for a microbe-mediate transition in developmental life-history depends on the ecological conditions of the host (*i.e.*, *B* parameter) as well as the manipulating capability of the microbe (*i.e.*, *V* parameter). These factors create a gradient in the sex ratio (A) needed to induce a transition in developmental life-history (*i.e.*, at 300 µm) as well as a gradient in the stable egg sizes that follow this transition (B), with the mean egg size for each combination presented here. No transition may also be observed when ecology heavily favors planktotrophy and the manipulating capability of the microbe is weak (gray, top left of A). When ecology favors larger egg sizes, only enhanced growth is needed for a transition to occur and no manipulation in sex ratio (light blue, bottom right of A). The microbe in this figure represents feminization and enhanced growth, but a nearly identical pattern is also observed for male killing and enhanced growth (Fig. 3).

**Fig. S8:**
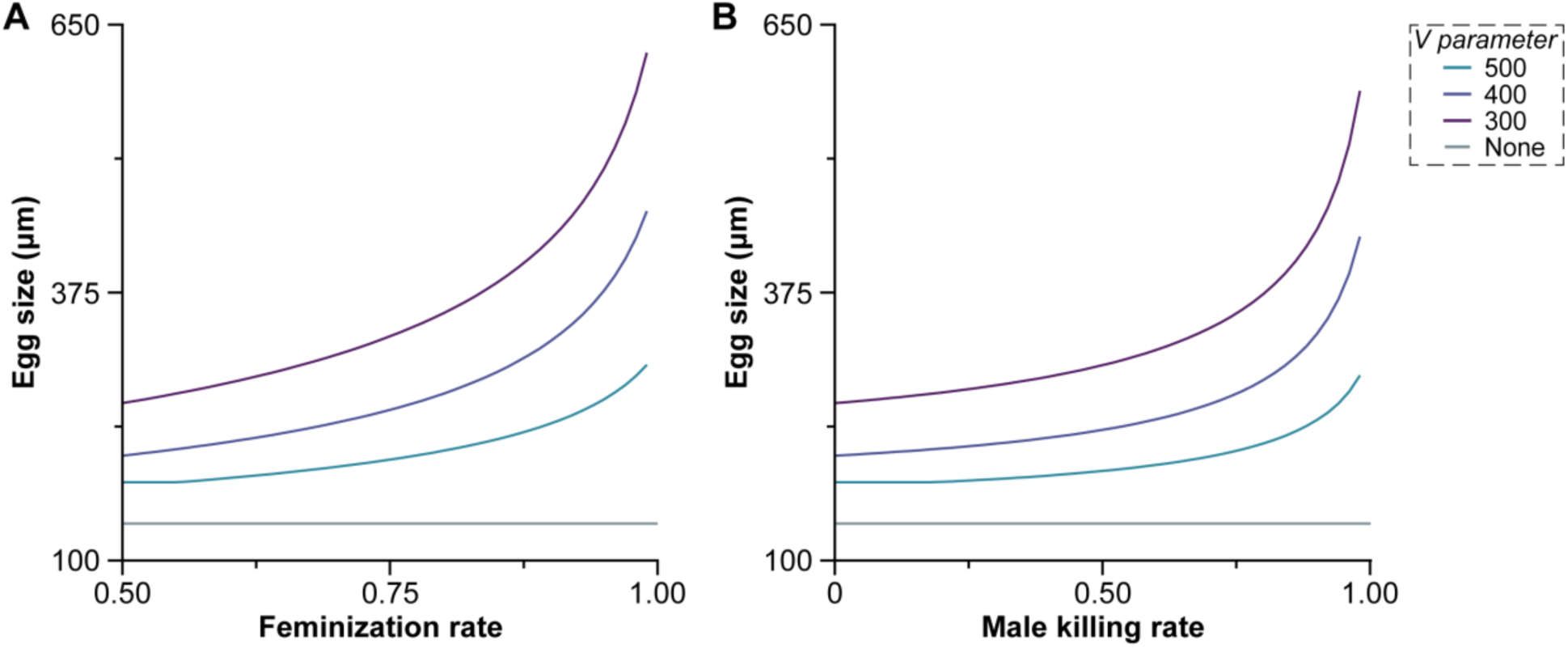
Egg size (μm in diameter) increases exponentially with feminization (left) and male killing (right) rates. Egg size increases at a given manipulation rate as *V* decreases. *B* parameter for this graph is set at 250. Grey line indicates evolutionary stable egg size for when there is no microbial association.

**Fig. S9:**
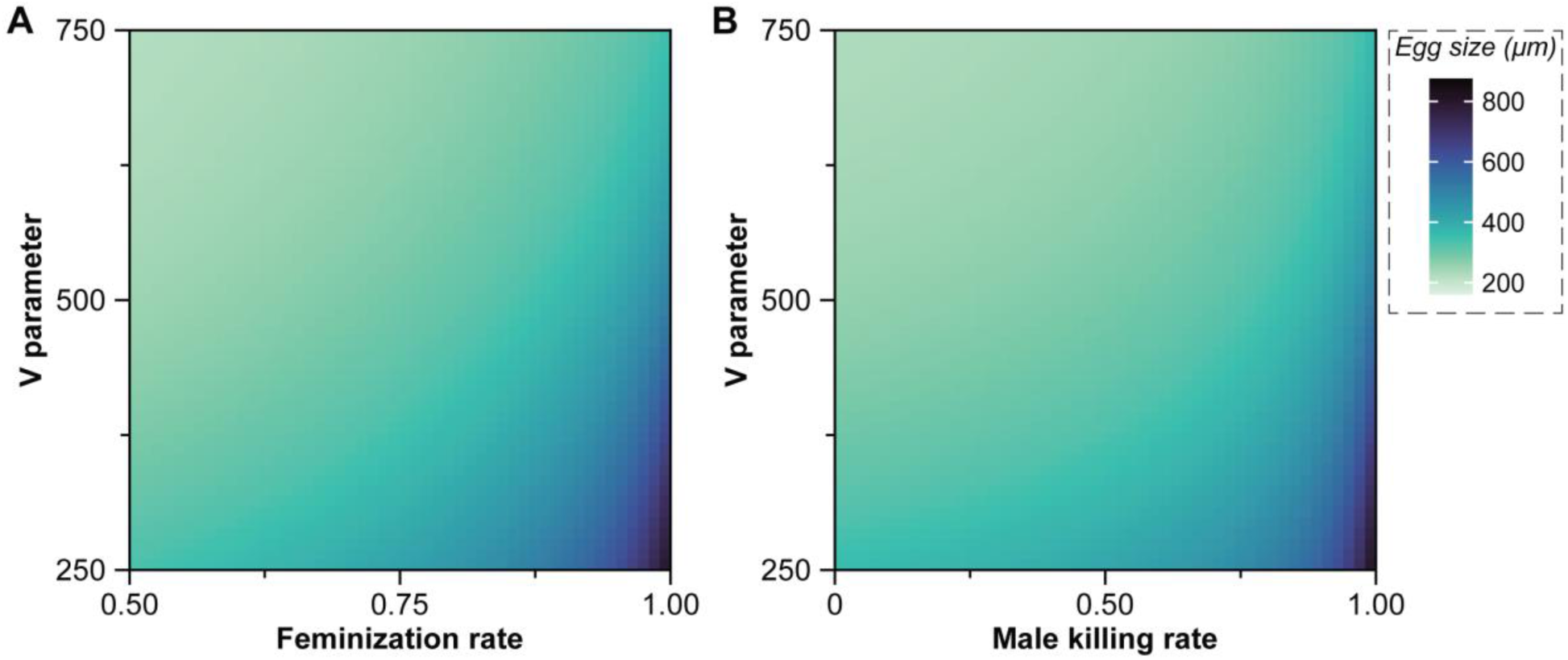
Stable maximum egg size increases with microbial manipulation and decreases with *V* parameter. This pattern is similar for a microbe that manipulates host reproduction by feminization (A, left) or male killing (B, right).

**Fig. S10:**
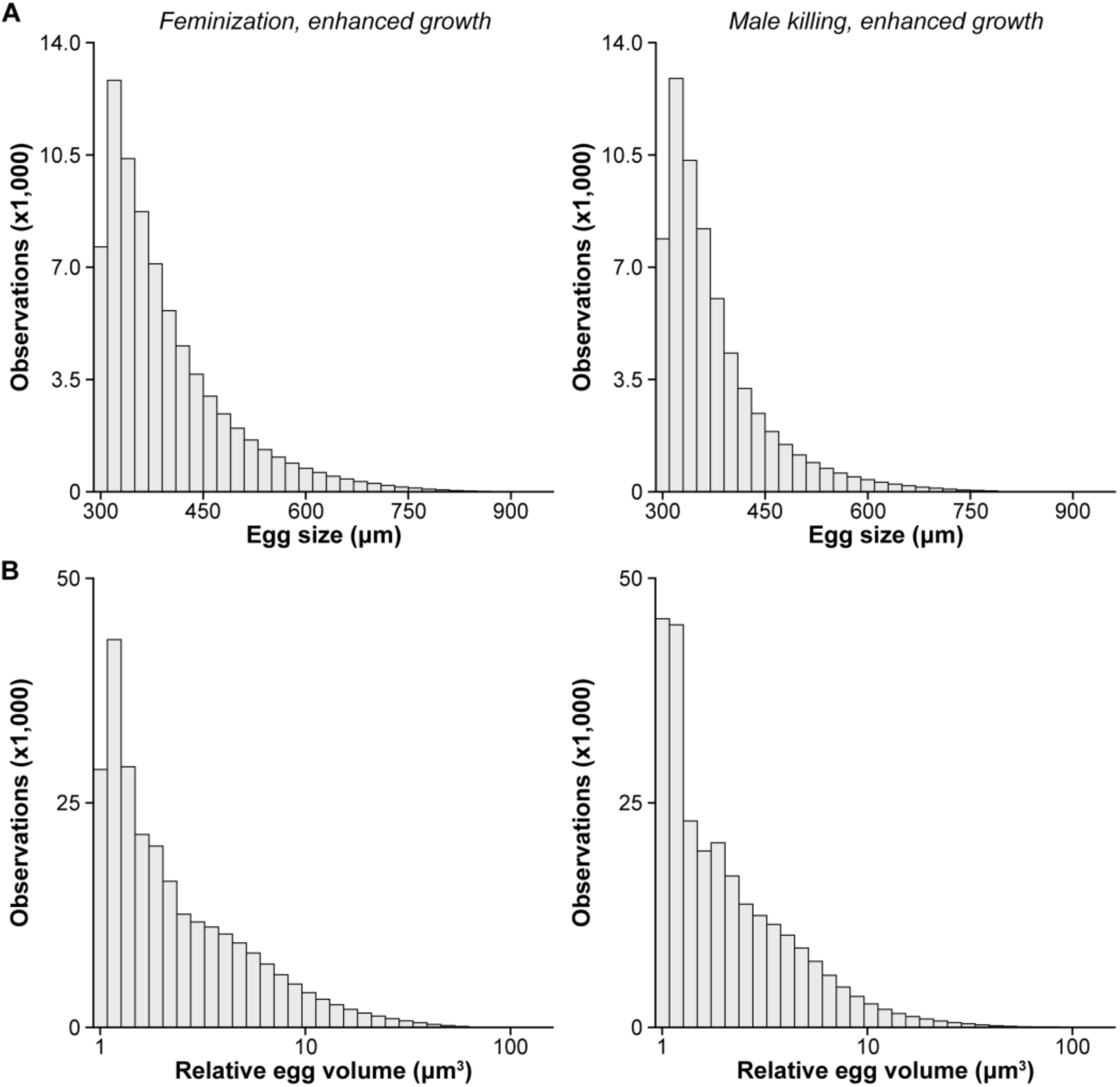
Stable egg sizes following a microbe-mediate transition in developmental life-history. Distribution of stable egg sizes (µm; A, top) and relative egg volume (µm^3^; B, bottom) following a transition in developmental life-history that was induced by a microbe that manipulates host reproduction by feminization (left) or male killing (right) and can compensate host fitness by supplementing host nutrition. These distributions represent the full range of stable egg sizes, as opposed to Fig. 3 that present the mean value for a given combination of the *B* and *V* parameters.

## Notes

### Competing Interest Statement

The authors have declared no competing interest.

